# Cellular Activities of SARS-CoV-2 Main Protease Inhibitors Reveal Their Unique Characteristics

**DOI:** 10.1101/2021.06.08.447613

**Authors:** Wenyue Cao, Chia-Chuan Dean Cho, Zhi Zachary Geng, Xinyu R. Ma, Robert Allen, Namir Shaabani, Erol C. Vatansever, Yugendar R. Alugubelli, Yuying Ma, William H. Ellenburg, Kai S. Yang, Yuchen Qiao, Henry Ji, Shiqing Xu, Wenshe Ray Liu

## Abstract

As an essential enzyme of SARS-CoV-2, the pathogen of COVID-19, main protease (M^Pro^) triggers acute toxicity to its human cell host, an effect that can be alleviated by an M^Pro^ inhibitor with cellular potency. By coupling this toxicity alleviation with the expression of an M^Pro^-eGFP fusion protein in a human cell host for straightforward characterization with fluorescent flow cytometry, we developed an effective method that allows bulk analysis of cellular potency of M^Pro^ inhibitors. In comparison to an antiviral assay in which M^Pro^ inhibitors may target host proteases or other processes in the SARS-CoV-2 life cycle to convene strong antiviral effects, this novel assay is more advantageous in providing precise cellular M^Pro^ inhibition information for assessment and optimization of M^Pro^ inhibitors. We used this assay to analyze 30 literature reported M^Pro^ inhibitors including MPI1-9 that were newly developed aldehyde-based reversible covalent inhibitors of M^Pro^, GC376 and 11a that are two investigational drugs undergoing clinical trials for the treatment of COVID-19 patients in United States, boceprevir, calpain inhibitor II, calpain inhibitor XII, ebselen, bepridil that is an antianginal drug with potent anti-SARS-CoV-2 activity, and chloroquine and hydroxychloroquine that were previously shown to inhibit M^Pro^. Our results showed that most inhibitors displayed cellular potency much weaker than their potency in direct inhibition of the enzyme. Many inhibitors exhibited weak or undetectable cellular potency up to 10 μM. On contrary to their strong antiviral effects, 11a, calpain inhibitor II, calpain XII, ebselen, and bepridil showed relatively weak to undetectable cellular M^Pro^ inhibition potency implicating their roles in interfering with key steps other than just the M^Pro^ catalysis in the SARS-CoV-2 life cycle to convene potent antiviral effects. characterization of these molecules on their antiviral mechanisms will likely reveal novel drug targets for COVID-19. Chloroquine and hydroxychloroquine showed close to undetectable cellular potency to inhibit M^Pro^. Kinetic recharacterization of these two compounds rules out their possibility as M^Pro^ inhibitors. Our results also revealed that MPI5, 6, 7, and 8 have high cellular and antiviral potency with both IC_50_ and EC_50_ values respectively below 1 μM. As the one with the highest cellular and antiviral potency among all tested compounds, MPI8 has a remarkable cellular M^Pro^ inhibition IC_50_ value of 31 nM that matches closely to its strong antiviral effect with an EC_50_ value of 30 nM. Given its strong cellular and antiviral potency, we cautiously suggest that MPI8 is ready for preclinical and clinical investigations for the treatment of COVID-19.

## INTRODUCTION

COVID-19 is an ongoing pandemic that has paralyzed much of the world. As of May 26^th^, 2021, the total confirmed infections have reached above 167 million and the total death toll has exceeded 3.4 million worldwide.^1^ The disease is ongoingly devastating countries including Brazil and India. With vaccines available for COVID-19, many countries have been conducting immunization campaigns hoping that herd immunity will be achieved when the majority of the population is vaccinated.^2^ Current COVID-19 vaccines are targeting the membrane Spike protein of SARS-CoV-2, the pathogen of COVID-19.^3^ Spike is a weakly conserved protein in a highly mutable RNA virus. Although SARS-CoV-2 shares overall 82% genome sequence identity with SARS-CoV, Spike has only 76% protein sequence identity shared between two origins.^4^ The highly mutable nature of Spike has also been corroborated by the continuous identification of new SARS-CoV-2 strains that have Spike mutations.^5^ The most notable are UK, South African, and currently Indian strains. Accumulated evidences have shown attenuated activity of developed vaccines against some newly emerged SARS-CoV-2 strains.^6^ Booster vaccines might be developed for new virus strains. However, the situation will likely turn into an incessant race between the emergence of new virus strains and the development of new vaccines. The focus on vaccine development and immunization that are preventative toward COVID-19 has largely obscured the development of targeted therapeutics that are direly needed for the treatment of patients with severe symptoms. By targeting a conserved gene in SARS-CoV-2, a small molecule medication can potentially turn more successful than a vaccine in containing the COVID-19 pandemic in both prevention and treatment since it is generally easier to manufacture, store, deliver, and administer a small molecule than a vaccine and the high conservativeness of the targeted gene will also make it hard for the virus to evade the small molecule.

One demonstrated drug target in SARS-CoV-2 is its main protease (M^Pro^).^7, 8^ Unlike Spike that is highly mutable, M^Pro^ is highly conserved. Its 96% protein sequence identity shared between SARS-CoV and SARS-CoV-2 is much higher than the overall 82% genome sequence identify shared between the two viruses.^3^ Much work has also been done in the development of M^Pro^ inhibitors.^9–11^ A general strategy that most researchers have been following in the development of M^Pro^ inhibitors is to synthesize an active site inhibitor, test its enzymatic inhibition, and then carry out its crystallographic and antiviral analysis to obtain information for next round optimization. For most medicinal chemists, the bottleneck in this drug discovery process is the antiviral assay that requires the use of a BSL3 facility and is often not accessible. The antiviral assay itself may also lead to misleading results about the real mechanism of an M^Pro^ inhibitor. The life cycle of SARS-CoV-2 (Figure 1A) requires a number of proteases that are from either the host or the virus itself. It has been shown that transmembrane protease serine 2 (TMPRSS2) serves a critical function to prime Spike for interactions with the human cell host receptor ACE2 during the virus entry process.^12^ After SARS-CoV-2 is internalized into an endosome, cathepsin L (CtsL) potentiates its membrane fusion with the endosome for the release of the virus RNA genome into the host cytosol.^13^ Other cathepsin proteins such as cathepsin B (CtsB) have also been suggested serving a role in the SARS-CoV-2 entry.^14^ After the SARS-CoV-2 genomic RNA is released into the host cytosol, it is translated by the host ribosome to form two large polypeptides, ORF1a and ORF1ab. The processing of OFR1a and ORF1ab to 15 mature nonstructural proteins (nsps) requires proteolytic functions of two internally coded protease fragments, nsp3 and nsp5 that are also called papain-like protease (PL^Pro^) and main protease (M^Pro^) respectively. Some nsps package into an RNA replicase complex that replicates both genomic and subgenomic RNAs. Translation of subgenomic RNAs leads to essential structural proteins for packaging new virions. Furin is a host protease that can hydrolyze Spike to prime it for new virion packaging and release.^15^ Based on our current understanding of SARS-CoV-2 pathogenesis and replication, there are at least three host and two viral proteases serving critical roles in the SARS-CoV-2 life cycle. Inhibition of any of these enzymes will potentiate a strong antiviral effect. Catalytic similarity between these enzymes also makes it likely that a developed small molecule is unselective toward these enzymes. M^Pro^, PL^Pro^, CtsB, and CtsL are cysteine proteases with a similar catalytic mechanism. TMPRSS2 and furin are serine proteases. Although serine proteases are mechanistically different from cysteine proteases, many currently developed M^Pro^ inhibitors have covalent warheads such as aldehyde and ketone making them prone to form covalent adducts with TMPRSS2 and furin as well to exert potent inhibition.^16, 17^ All these proteases are also localized in different parts of the host cell. Their inhibition requires different characteristics in their inhibitors such as cellular permeability and pH sensitivity. A simple antiviral assay of a developed M^Pro^ inhibitor will likely lead to a positive result that reflects inhibition not necessarily of M^Pro^ and therefore causes misunderstanding that can be detrimental to further rounds of lead optimization. Therefore, an assay system that directly reflects M^Pro^ inhibition in the host cell is critical for both assessment and optimization of M^Pro^ inhibitors. In the current work, we will describe such a system and its application in revealing unique characteristics of a number of developed and repurposed M^Pro^ inhibitors.

**Figure 1:**
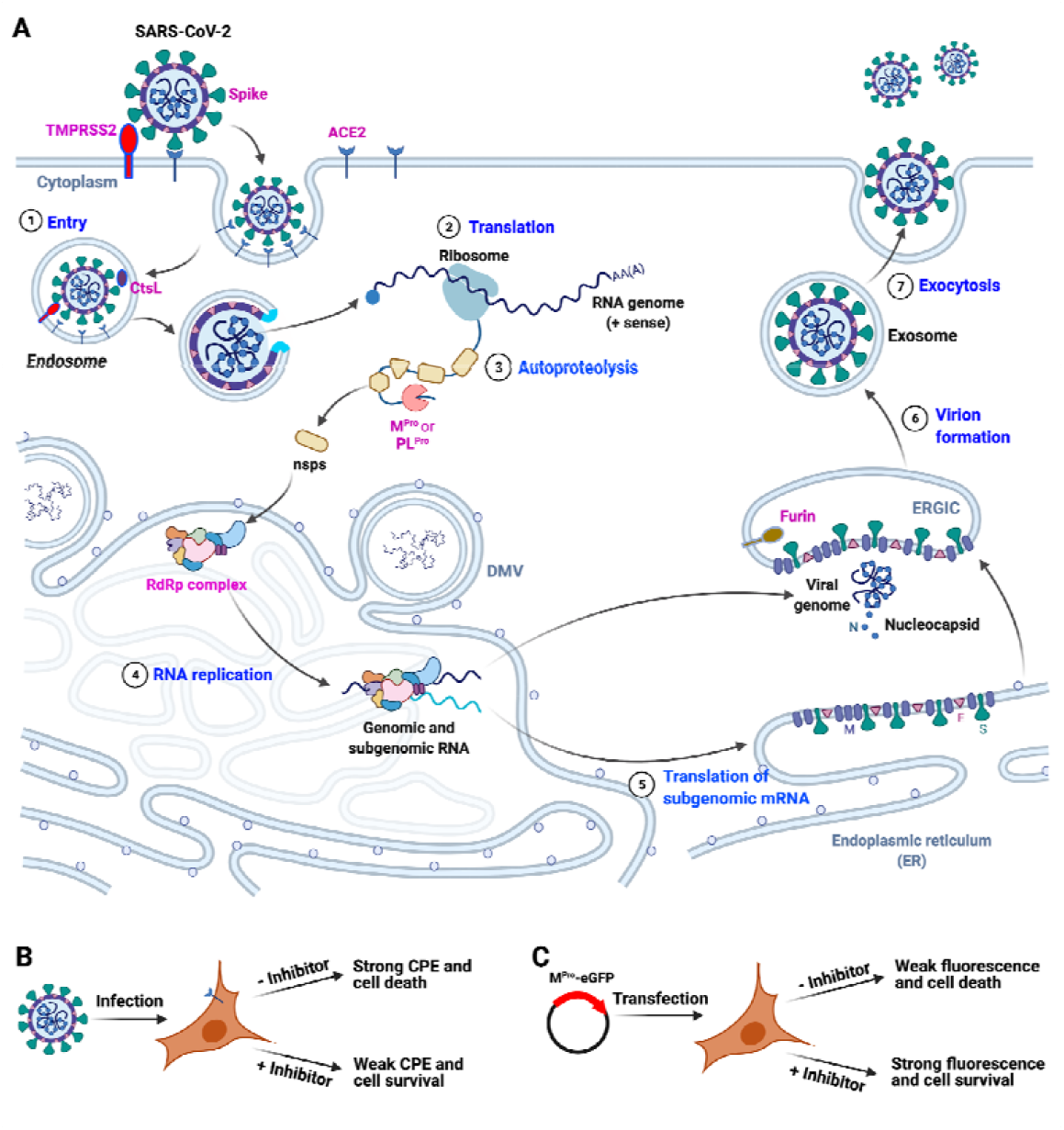
The life cycle of SARS-CoV-2 and two assays for M^Pro^-targeting antivirals. (**A**) A cartoon diagram illustrating the life cycle of SARS-CoV-2. Seven sequential steps are labeled in blue. Proteins that are labeled in pink are targets for the development of antivirals. TMPRSS2, CtsL and furin are three host proteases that prime Spike for viral entry and new virion packaging. ACE2: angiotensin-converting enzyme 2; TMPRSS2: transmembrane protease serine 2; CtsL: cathepsin L; M^Pro^: main protease; PL^Pro^: papain-like protease; RdRp: RNA-dependent RNA polymerase; nsp: nonstructural protein. (**B**) An antiviral assay based on the inhibition of virus infection-triggered cytopathogenic effect (CPE) and cell death. (**C**) An antiviral assay based on the inhibition of M^Pro^-induced apoptosis in host cells and the fluorescence of the expressed M^Pro^-eGFP fusion protein.

## RESULTS

### The rationale and the establishment of a cellular M^Pro^ inhibition assay for MPI8

A typical antiviral assay for SARS-CoV-2 is its triggering of strong cytopathogenic effect (CPE) in host cells leading to death that can be quantified by counting formed viral plaques (Figure 1B). An M^Pro^ inhibitor with high cellular potency will suppress this strong CPE leading to host cell survival. A good cellular M^Pro^ inhibition assay will need to mimic this CPE suppression process to a large extent. Our original design for a cellular M^Pro^ inhibition assay was to express M^Pro^ in host cells that is fused with a *N*-terminal cyan fluorescent protein (CFP) and a *C*-terminal yellow fluorescent protein (YFP) and test the inhibition of autocleavage of this fusion protein in the presence of an inhibitor. M^Pro^ natively cuts off its fused protein at the *C*-terminus. We put an M^Pro^ digestion site between CFP and M^Pro^ for its cleavage as well. CFP and YFP form a Förster resonance energy transfer (FRET) pair.^18^ Without an inhibitor, both CFP and YFP will be cleaved from the fusion protein in host cells leading to no FRET signal. In the presence of a potent inhibitor, the fusion protein will be intact in host cells leading to strong FRET signals. However, transfection of 293T cells with pECFP-M^Pro^-EYFP (SI Appendix, Fig. S1), a plasmid containing a gene coding the CFP-M^Pro^-YFP fusion protein led to death of most transfected cells. Repeating this transfection process all led to the exact same result. It is evident that M^Pro^ can exert acute toxicity to its human cell host. The same observation has been made by others as well.^19^ MPI8 is an M^Pro^ inhibitor that our lab developed previously.^16^ Antiviral analysis indicated that MPI8 has potency to totally suppress SARS-CoV-2-induced CPE in ACE2^+^ A549 cells with a concentration around 0.2 μM. Given its approved antiviral potency, we used MPI8 as a positive control molecule for the analysis of cellular M^Pro^ inhibition. To alleviate the toxicity that was induced by the expression of CFP-M^Pro^-YFP, we cultured 293T cells that were transfected with pECFP-M^Pro^-EYFP in media containing 10 μM MPI8. The presence of MPI8 reduced death of transfected cells sharply. Interestingly the overall expressed fusion protein was also significantly improved, showing much enhanced, directly detected yellow fluorescence from YFP (SI Appendix, Fig. S2). This positive correlation between the expression of CFP-M^Pro^-YFP and the survival of transfected cells is likely due to the shutting-down of translation by active M^Pro^. In comparison to the measurement of inhibitor-induced FRET signal increase in CFP-M^Pro^-YFP, the measurement of cellular survival-correlated fluorescence improvement from CFP-M^Pro^-YFP in the presence of an inhibitor mimics the suppression of virus-induced CPE by an inhibitor in a real antiviral assay more since the antiviral assay is also based on the host cell survival. Therefore, we decided to adopt this new way for the analysis of cellular potency of M^Pro^ inhibitors.

Since a FRET system is not necessary for cellular potency analysis of M^Pro^ inhibitors, we modified our plasmid to express an M^Pro^-eGFP fusion protein (Figure 1C) in host cells that can be easily analyzed using fluorescent flow cytometry. The expression of M^Pro^-eGFP in host cells will trigger cell death and weak fluorescence. This process will be reversed by adding a potent inhibitor with cellular activity. In order to use eGFP fluorescence to accurately represent expressed M^Pro^, we introduced a Q306G mutation in M^Pro^ to abolish its cleavage of the *C*-terminal eGFP. M^Pro^ requires a free *N*-terminal serine for strong activity. To achieve this, we built two constructs as shown in Figure 2A and SI Appendix, Fig S3. The first construct pLVX-M^Pro^-eGFP-1 encodes M^Pro^-eGFP with a *N*-terminal methionine that relies on host methionine aminopeptidases for its cleavage. The second construct pLVX-M^Pro^-eGFP-2 encodes M^Pro^-eGFP containing a short *N*-terminal peptide that has an M^Pro^ cleavage site at the end for its autocatalytic release. Transfection of 293T cells with two constructs showed that pLVX-M^Pro^-eGFP-2 led to more potent toxicity to cells and this toxicity was effectively suppressed when we provided 10 μM MPI8 in the growth media (Figure 2B). Therefore, we selected pLVX-M^Pro^-eGFP-2 for all our following studies. To demonstrate that cellular fluorescence is positively correlated to the concentration of provided MPI8, we transfected 293T cells with pLVX-M^Pro^-eGFP-2, grew transfected cells in the presence of four MPI8 concentrations (0, 20, 40, and 160 nM) for 72 h, and then sorted cells using fluorescent flow cytometry (Figure 2C). Both the number and intensity of fluorescent cells (FL1-A signal > 1 ×10^6^) were positively dependent on the provided MPI8 concentration, indicating the feasibility of using the system to characterize cellular potency of an M^Pro^ inhibitor. To demonstrate this feasibility, we transiently transfected 293T cells with pLVX-M^Pro^-eGFP-2 and grew transfected cells in the presence of a cascade of MPI8 concentrations that started from 10 μM and descended 5 folds consecutively. After 72 h, we sorted cells according to their eGFP fluorescent intensity. Cells with FL1-A signal above 1 × 10^6^ were analyzed. We built a METLAB script to calculate average eGFP fluorescent intensity of all analyzed cells and plotted average eGFP fluorescent intensity against the MPI8 concentration as shown in Figure 3D. The data showed obvious MPI8-induced saturation of M^Pro^-eGFP expression and fit nicely to a three-parameter dose dependent inhibition mechanism in Prism 9 for IC_50_ determination. The determined cellular M^Pro^ inhibition IC_50_ value of MPI8 is 31 nM. As presented later, an antiviral assay in Vero E6 cells showed an EC_50_ value of 30 nM for MPI8 in inhibiting SARS-CoV-2. This high similarity between cellular M^pro^ inhibition IC_50_ and antiviral EC_50_ values of MPI8 validates that cellular M^Pro^ inhibition potency of an inhibitor represents closely its antiviral potency through M^Pro^ inhibition.

**Figure 2:**
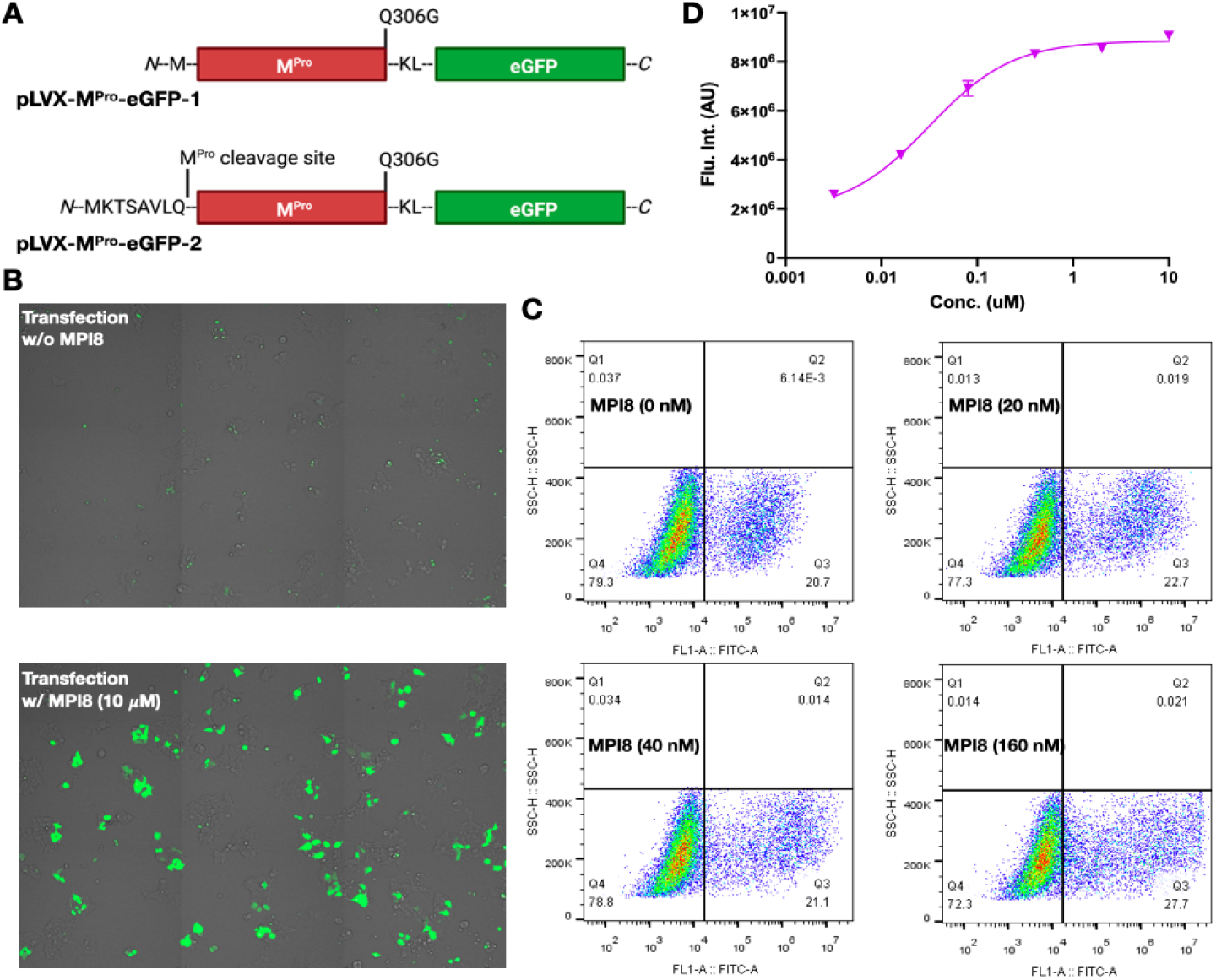
The validation of transiently expressed M^Pro^ and its cellular toxicity for the analysis of cellular potency of M^Pro^ inhibitors. (**A**) The design of two M^Pro^-eGFP fusions. The first design requires *N*-terminal methionine processing for M^Pro^ activation, and the second design relies on autocleavage at the *N*-terminal M^Pro^ cleavage site for M^Pro^ activation. (**B**) 293T cells transiently transfected with pLVX-M^Pro^-eGFP-2 and grown in the absence or presence of 10 μM MPI8. Cells grown in the absence of MPI8 had a low level of M^Pro^-eGFP expression and a high level of cell death but cells grown in the presence of MPI8 had a high level of M^Pro^-eGFP expression and a low level of cell death. (**C**) 293T cells that were transiently transfected with pLVX-M^Pro^-eGFP-2 expressed M^Pro^-eGFP correlated with the concentration of MPI8 in the growth media. (**D**) The cellular IC_50_ determination of MPI8. 293T cells were transfected with pLVX-M^Pro^-eGFP-2 and grown in the presence of different concentrations of MPI8 for 72 h before their sorting using flow cytometry. Average fluorescent intensity for cells with FL1-A signal higher than 2 × 10^6^ was determined and used to plot against the MPI8 concentration. Data were fit to the three-parameter dose dependent inhibition mechanism to determine the cellular IC_50_ value.

**Figure 3:**
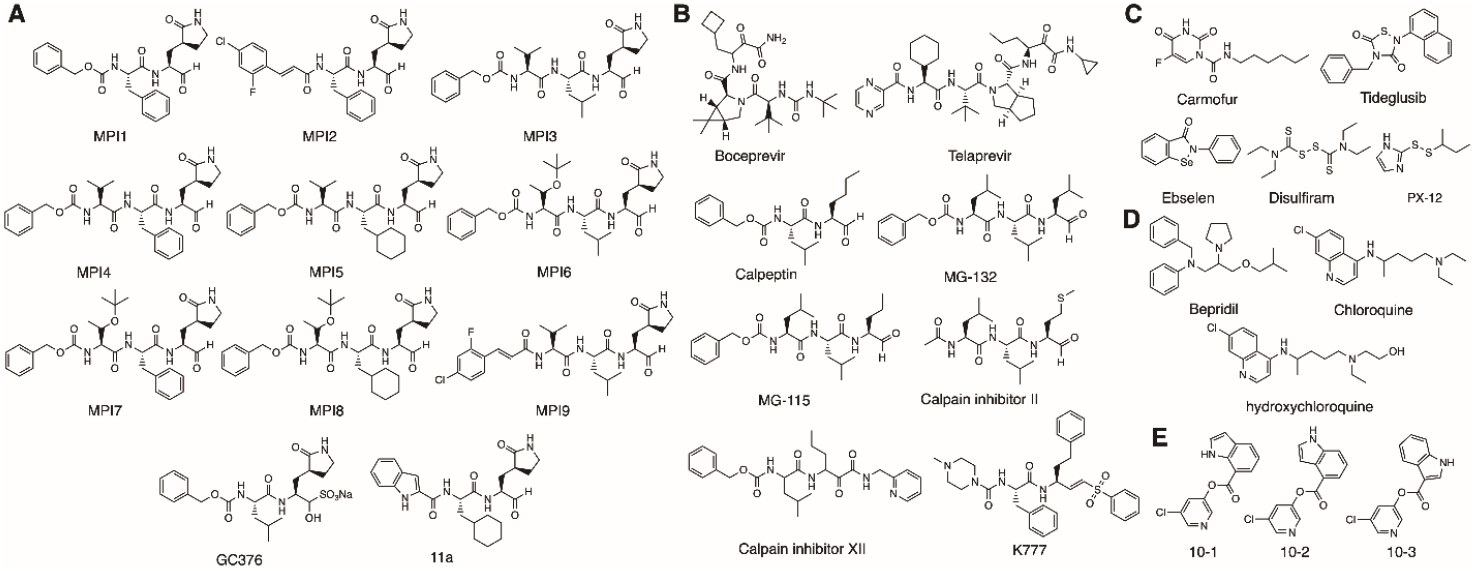
The structures of inhibitors that were investigated in their cellular inhibition of M^Pro^. (**A**) Reversible covalent inhibitors designed for M^Pro^. (**B**) Investigational covalent inhibitors that were developed for other targets. (**C**) Inhibitors that were identified via high-throughput screening. (**D**) FDA-approved medications that have been explored as M^Pro^ inhibitors. (E) Diaryl esters that have high potency to inhibit M^Pro^.

Since MPI8 is highly effective in inhibiting M^Pro^ in cells, we used it in combination with pLVX-M^Pro^-eGFP-2 to make stable 293T cells that continuously expressed M^Pro^-eGFP. Using this stable cell line, we characterized M^Pro^-induced apoptosis that was detected by anti-annexin. After we withdrew MPI8 from the growth media that we used to culture stable cells, strong apoptotic effect started to show after 24 h and continued to increase (SI Appendix, Fig. S4). Since MPI8 is a reversible covalent inhibitor, the relatively long incubation time for the observation of apoptosis is likely due to its slow release from the M^Pro^ active site. Due to concerns about residual MPI8 and its potential slow release from M^Pro^ in stable cells, we chose to perform cellular potency characterization of all M^Pro^ inhibitors by doing transient transfection of 293T cells and then growth in the presence of different inhibitor concentrations.

### MPI1-7, MPI9, GC376, and 11a

MPI8 was one of 9 β-(*S*-2-oxopyrrolidin-3-yl)-alaninal (Opal)-based, reversible covalent M^Pro^ inhibitors MPI1-9 we previously developed (Figure 3A).^16^ GC376 is a prodrug that dissociates quickly in water to release its Opal component.^20^ 11a is another Opal-based, reversible covalent M^Pro^ inhibitor that was developed in 2020.^9^ All 11 compounds showed high potency in inhibiting M^Pro^ in an enzymatic assay.^16^ Besides MPI8, we went on to test cellular potency of all other 10 Opal inhibitors in their cellular inhibition of M^Pro^ as well by following the exact same procedure that we did for MPI8. As shown in Figure 4A, all tested Opal inhibitors promoted cell survival and the expression of M^Pro^-eGFP significantly at 10 μM. However, data collected at different concentrations showed that only three inhibitors, MPI5, 6, and 7 induced saturation of M^Pro^-eGFP expression at or below 10 μM. Determined IC_50_ values for MPI5, 6, and 7 are 0.66, 0.12, and 0.19 μM, respectively (Table 1). Based on collected data, MPI2-4, MPI9, GC376, and 11a have IC_50_ values higher than 2 μM and MPI1 that displayed the lowest inhibition of M^Pro^ at 10 μM among all Opal inhibitors has an IC_50_ value higher than 10 μM.

**Figure 4:**
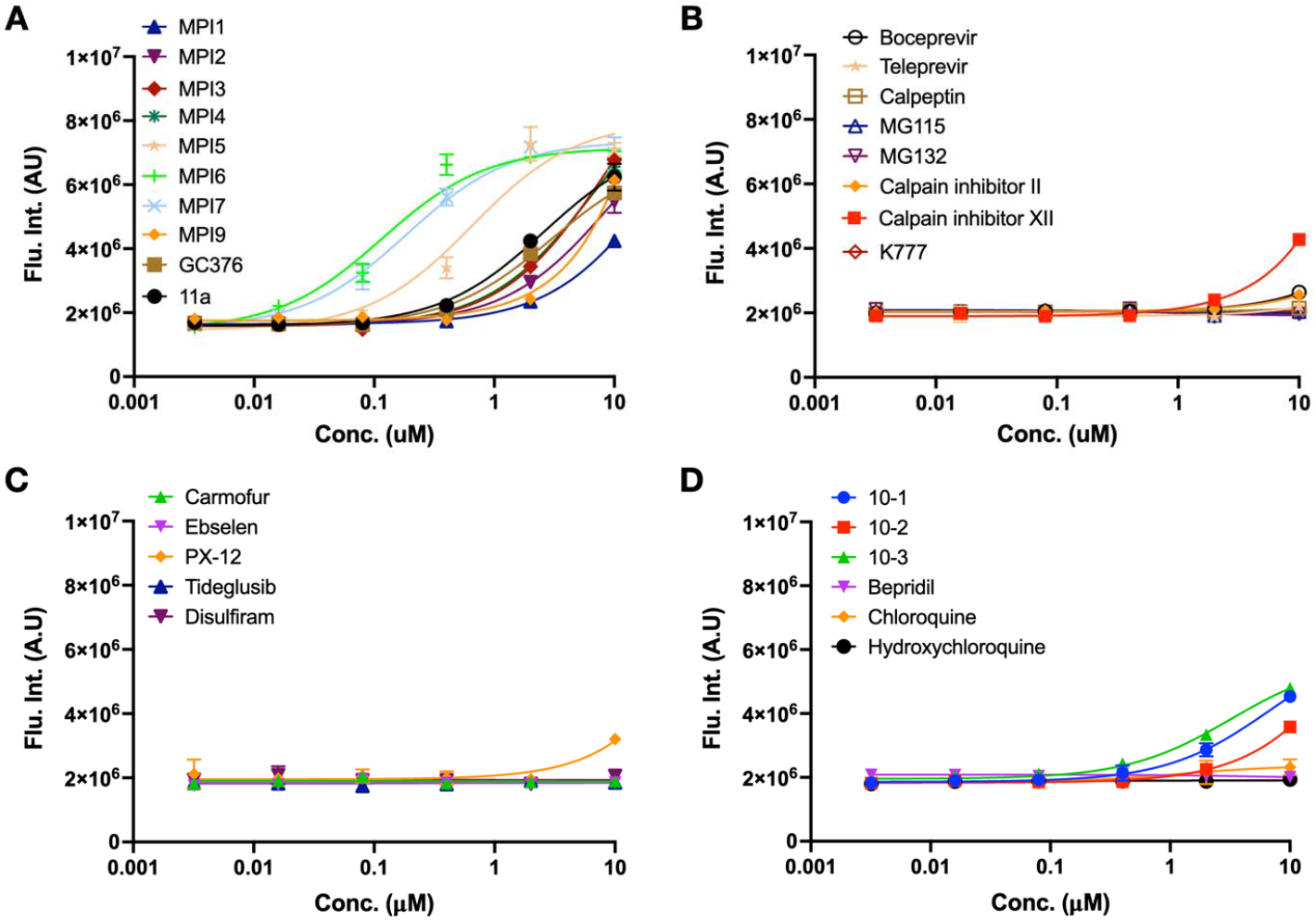
Cellular potency of literature reported M^Pro^ inhibitors. K777 is included as a potential M^Pro^ inhibitor.

**Table 1:**
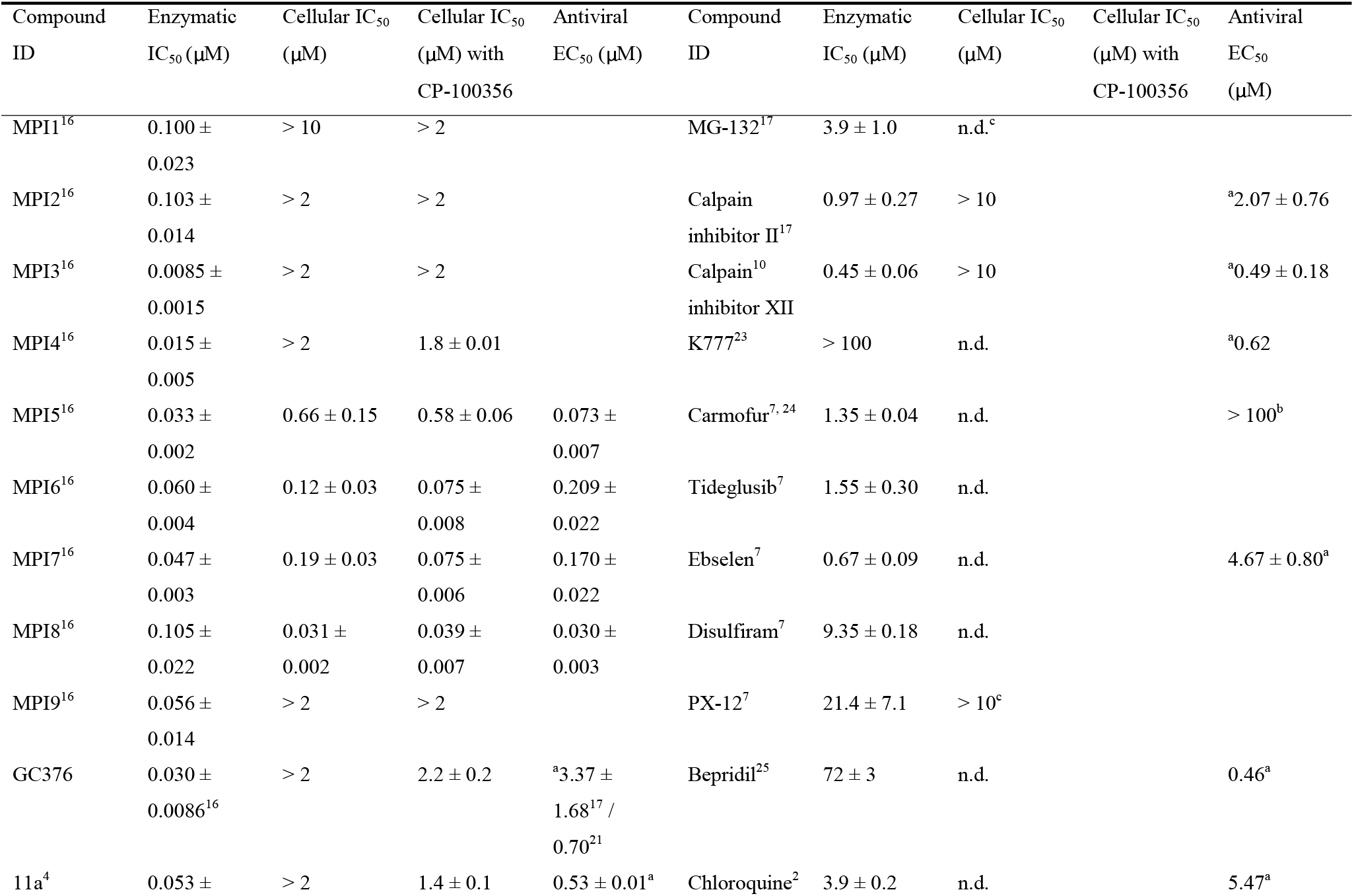
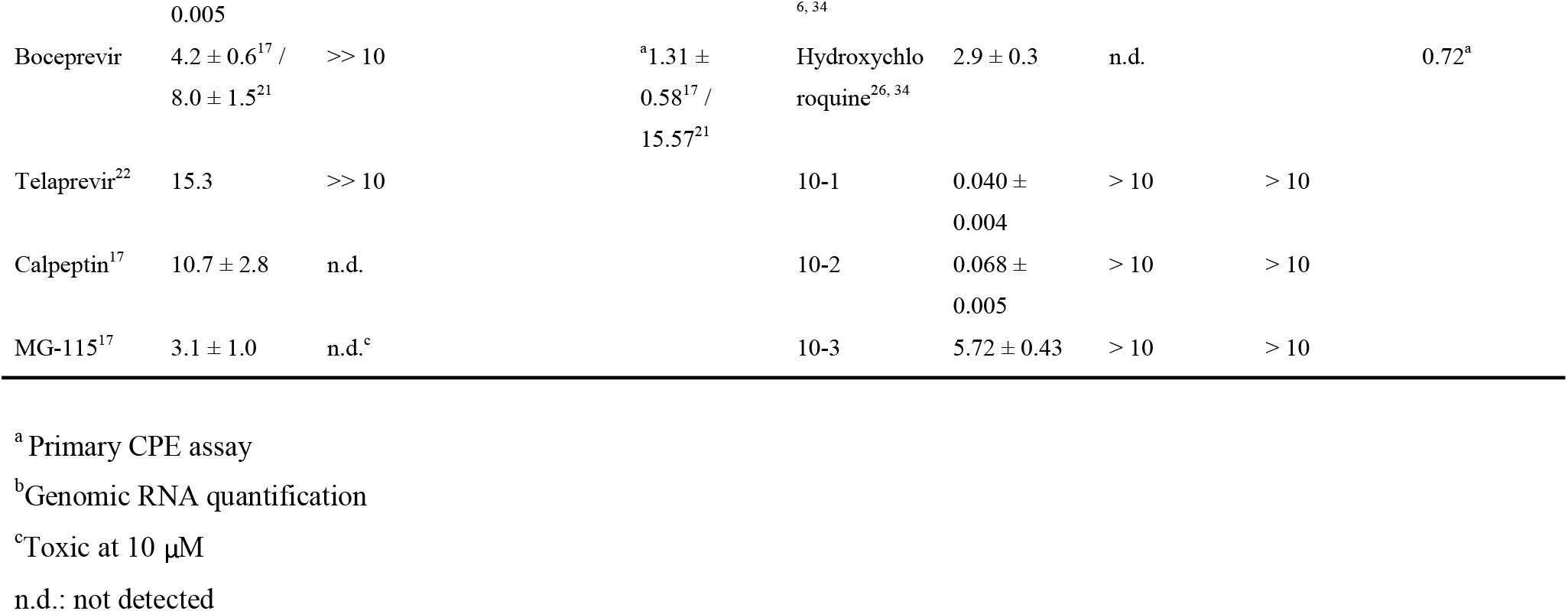
Determined enzymatic and cellular IC_50_ values in inhibiting SARS-CoV-2 M^Pro^ for different inhibitors

### Boceprevir, telaprevir, calpeptin, MG-132, MG-115, calpain inhibitor II, calpain inhibitor XII, and K777

Drug repurposing research has led to the identification of a number of both FDA-approved and investigational medications as M^Pro^ inhibitors. These include boceprevir, telaprevir, and calpain inhibitor XII that have an α-ketoamide moiety for the formation of a reversible covalent adduct and calpeptin, MG-132, and calpain inhibitor II that has an aldehyde for a reversible covalent interaction with the M^Pro^ active site cysteine.^17, 21, 22^ Some of these compounds display potency in inhibiting SARS-CoV-2 replication in host cells as well. We went on to characterize cellular potency of these inhibitors using our developed cellular assay. K777 is a known CtsL inhibitor with high potency in inhibiting SARS-CoV-2 replication in human cell host.^23^ It has a vinylsulfonate moiety. Due to its propensity to form a permanent covalent adduct with the M^Pro^ active site cysteine, we tested its cellular potency in inhibiting M^Pro^. As shown in Figure 4B, calpeptin, MG115, MG132, telaprevir, and K777 displayed close to undetectable cellular inhibition of M^Pro^ up to 10 μM, boceprevir and calpain inhibitor II displayed close to undetectable cellular inhibition of M^Pro^ up to 2 μM and very weak cellular inhibition of M^Pro^ at 10 μM, and calpeptin XII exhibited highest cellular inhibition of M^Pro^ among this group of inhibitors but its inhibition activity is low with an estimated IC_50_ value higher than 10 μM.

### Carmofur, tideglusib, ebselen, disulfiram, and PX-12

Drug repurposing research has also shown that carmofur, tideglusib, ebselen, disulfiram, and PX-12 can potently inhibit M^Pro^.^7^ Carmofur is an antineoplastic agent that generates a permanent thiocarbamate covalent adduct with the M^Pro^ active site cysteine.^24^ All other four compounds are redox active for covalent conjugation with the M^Pro^ active site cysteine. We applied our cellular potency assay to these drugs as well. As shown in Figure 4C, except PX-12 that weakly inhibited M^Pro^ in cells that led to weak promotion of cell survival and M^Pro^-eGFP expression at 10 μM, the other four compounds showed undetectable cellular inhibition of M^Pro^ at all tested inhibitor concentrations.

### Bepridil, chloroquine, and hydroxychloroquine

Using computational docking analysis in combination with experimental examination to guide drug repurposing for COVID-19, we previously showed that bepridil, an antianginal drug inhibited M^Pro^ and had high potency in inhibiting SARS-CoV-2 replication in host cells.^25^ To provide a full picture for understanding the mechanism of bepridil in inhibiting SARS-CoV-2, we used our cellular M^Pro^ inhibition assay to study bepridil as well. As shown in Figure 4D, bepridil displayed very weak inhibition of M^Pro^ in cells up to 10 μM. A previous publication reported that chloroquine and hydroxychloroquine are potent inhibitors of M^Pro^.^26^ We tested these two drugs in inhibiting M^Pro^ in cells. At all tested concentrations, both drugs displayed close to undetectable promotion of M^Pro^-eGFP expression indicating very low M^Pro^ inhibition from both drugs in cells. Using both a commercial and home-made substrate, we recharacterized M^Pro^ enzymatic inhibition by chloroquine and hydroxychloroquine. Our data (SI Appendix, Figure S5) show that M^Pro^ retains 84% activity at 16 μM chloroquine in an enzyme activity assay. In the same assay, hydroxychloroquine do not inhibit M^Pro^ up to 16 μM.

### Diarylesters 10-1, 10-2, and 10-3

Benzotriazole esters were contaminants in a peptide library that were accidentally discovered as potent inhibitors of SARS-CoV M^Pro^.^27, 28^ Based on their inhibition mechanism of SARS-CoV M^Pro^, a number of diarylesters were developed later as potent SARS-CoV M^Pro^ inhibitors.^29, 30^ To show whether similar compounds will also inhibit M^Pro^ of SARS-CoV-2, we synthesized diarylesters 10-1, 10-2, and 10-3. Characterization of three compounds using an enzymatic inhibition assay resulted enzymatic inhibition IC_50_ values as 0.067, 0.038, and 7.6 μM for 10-1, 10-2, and 10-3, respectively (SI Appendix, Fig. S6). Using our cellular inhibition assay, we characterized all three compounds as well. As shown in Figure 4D, all three compounds display observable potency in inhibiting M^Pro^ to promote M^Pro^-eGFP expression at 2 and 10 μM. Their cellular M^Pro^ inhibition IC_50_ values are estimated above 10 μM.

### The effect of CP-100356 on cellular potency of peptide-based M^Pro^ inhibitors

CP-100356 is a high affinity inhibitor of multi-drug resistance protein (Mdr-1/gp), a protypical ABC transport that exports toxic substances from the inside of cells. A previous report showed that CP-100356 enhanced antiviral potency of M^Pro^ inhibitors dramatically.^31^ To investigate whether CP-100356 improves cellular M^Pro^ inhibition potency of Opal inhibitors, we recharacterized MPI1-9, GC376, and 11a using our cellular M^Pro^ inhibition assay in the presence of 0.5 μM CP-100356 (Figure 5). Except MPI8 that showed an inhibition curve in the presence of CP-100356 very similar to that in the absence of CP-100356 and had a determined IC_50_ value as 39 nM, all other Opal inhibitors displayed a better cellular M^Pro^ inhibition curve. MPI5 and MPI6 have IC_50_ values (580 and 75 nM respectively) in the presence of CP-100356 slightly lower than that in the absence of CP-100356. The highest cellular potency improvement that we observed among all compounds was for MPI7. It displayed an IC_50_ value (75 nM) in the presence of CP-100356 60% lower than that in the presence of CP-100356. The cellular potency improvement for MPI4, GC376, and 11a in the presence of CP-100356 also led to their IC_50_ values able to be determined as 1.8, 2.2, and 1.4 μM, respectively. We did a similar test with 10-1, 10-2, and 10-3. Providing CP-100356 did not significantly change cellular M^Pro^ inhibition for all three compounds at all tested concentrations.

**Figure 5:**
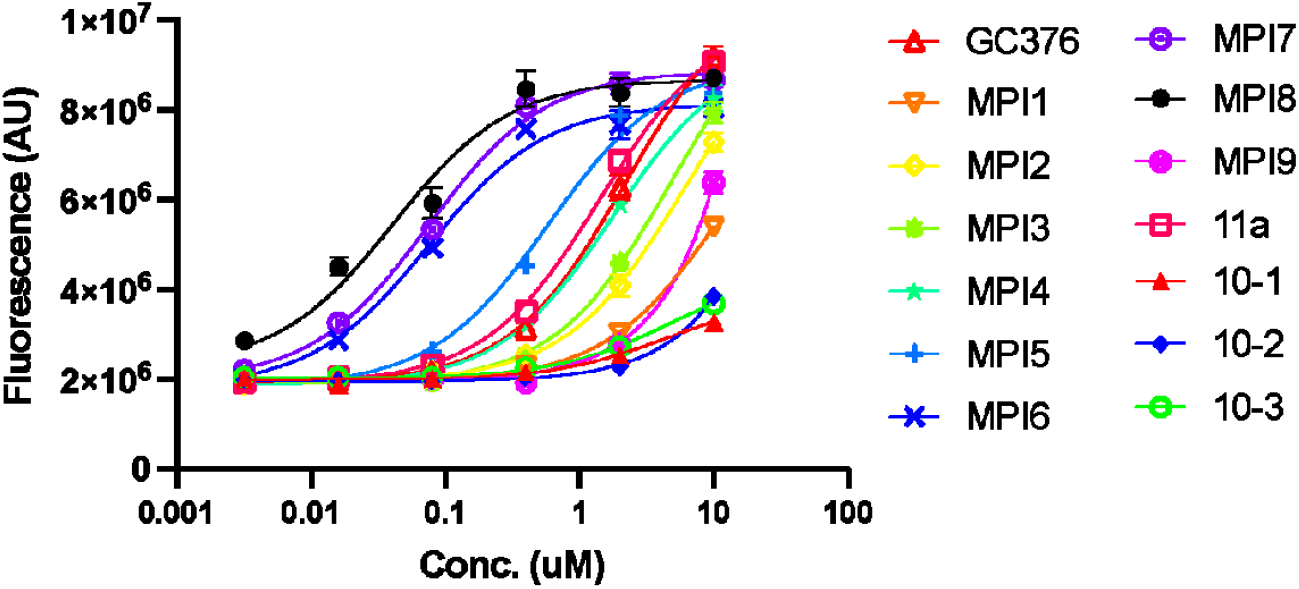
Cellular potency of selected compounds in their inhibition of M^Pro^ in the presence of 0.5 μM CP-100356.

### The determination of antiviral EC_50_ values for MPI5-8

Our previous antiviral assay for Opal inhibitors were based on on-off observation of CPE in Vero E6 and ACE2^+^ A549 cells. To quantify antiviral EC_50_ values of MPI5-8, we conducted plaque reduction neutralization tests of SARS-CoV-2 in Vero E6 cells in the presence of MPI5-8. we infected Vero E6 cells with SARS-CoV-2, grew infected cells in the presence of different concentrations of each inhibitor for 3 days, and then quantified SARS-CoV-2 plaque reduction. Based on SARS-CoV-2 plaque reduction in the presence of MPI5-8, we determined antiviral EC_50_ values for MPI5-8 as 73, 209, 170, and 30 nM, respectively (Figure 6).

**Figure 6:**
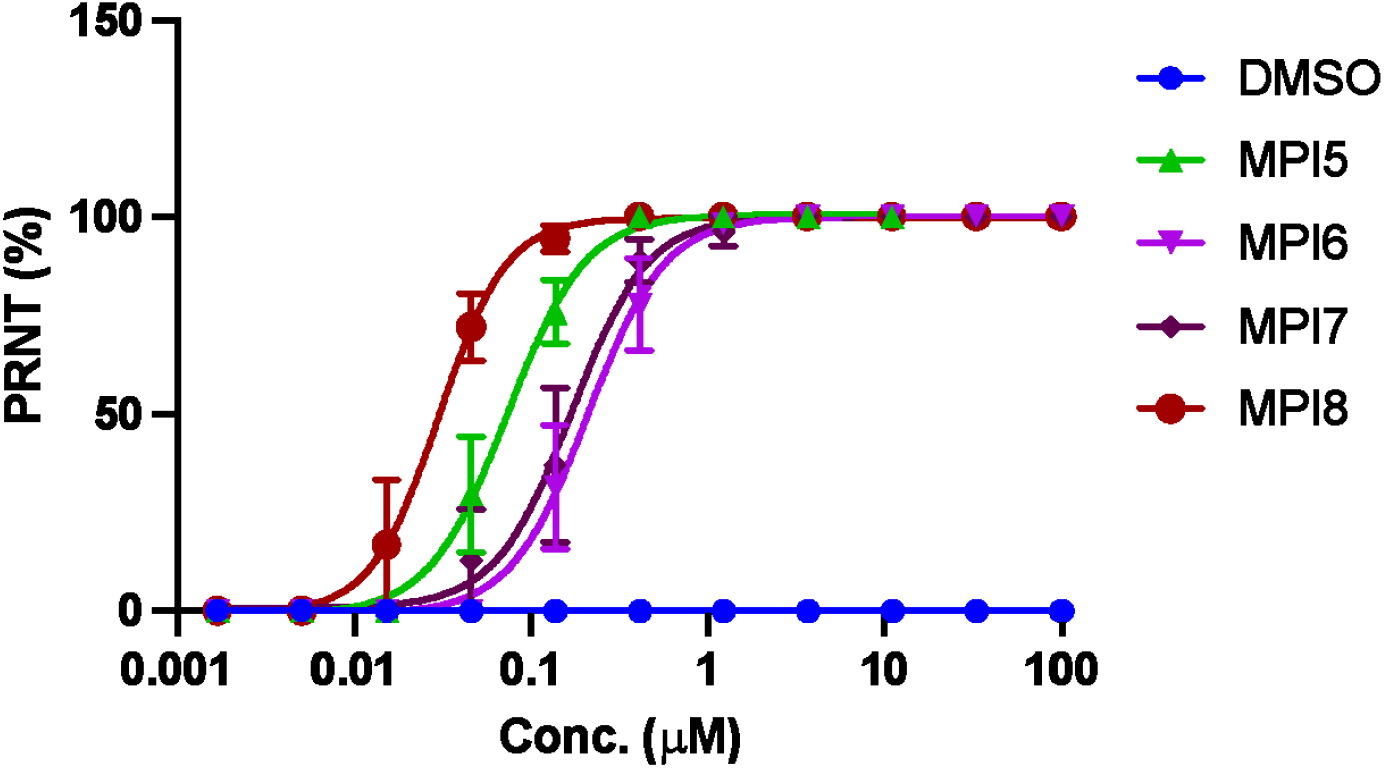
Plaque reduction neutralization tests (PRNTs) of MPI5-8 on their inhibition of SARS-CoV-2 in Vero E6 cells. DMSO was used as a negative control.

## DISCUSSION

The SARS-CoV-2 life cycle requires the involvement of proteases from both the virus and the human cell host. Given high similarity in catalytic mechanisms of these proteases, an inhibitor that is developed for M^Pro^ may also inhibit other proteases in the SARS-CoV-2 pathogenesis and replication pathway to exert an antiviral effect. Therefore, a direct antiviral assay is not optimal to reveal the real antiviral mechanism of an inhibitor and for its structureactivity relationship study for optimization. The strict requirement of a BSL3 facility to handle SARS-CoV-2 also prevents many research groups from conducting an antiviral assay in their labs and therefore causes delays in drug development. The antiviral assay itself is also complicated, lengthy, and difficult to turn high throughput. To resolve these issues, we developed a cellular M^Pro^ inhibition assay that can be easily characterized using fluorescent cell cytometry for bulk analysis of M^Pro^ inhibitors. We applied this assay to analyze 30 claimed M^Pro^ inhibitors and revealed unique features for a number of them.

MPI1-9 were previously developed as potent M^Pro^ inhibitors. All showed enzymatic IC_50_ values around or below 100 nM (Table 1). Among them MPI3 has the most enzymatic inhibition potency with an IC_50_ value of 8.5 nM. However, a CPE-based antiviral assay in Vero E6 cells showed that MPI3 weakly inhibited SARS-CoV-2.^16^ On the contrary, MPI8 that has an enzymatic IC_50_ value of 105 nM displayed the most potency in inhibiting SARS-CoV-2. A separate antiviral assay in ACE2^+^ A549 cells showed that MPI8 inhibited CPE from SARS-CoV-2 completely around 200 nM MPI8. Overall, the antiviral potency of MPI1-9 correlates well with their cellular M^Pro^ inhibition potency that we have detected using the new cellular assay. To see if our cellular potency results matched closely real antiviral effects, we quantified antiviral effects of MPI5-8 against SARS-CoV-2 in Vero E6 cells. Except MPI5 that displayed a 9-fold less antiviral EC_50_ value than its cellular IC_50_ value, our determined antiviral EC_50_ values for MPI6-8 closely matched their cellular M^Pro^ inhibition IC_50_ values validating the cellular inhibition assay in calibrating antiviral potency of M^pro^ inhibitors. Cellular potency for compounds determined by this new assay represents closely their antiviral potency through M^Pro^ inhibition. Our determined antiviral EC_50_ value for MPI8 is almost identical to its cellular M^Pro^ inhibition IC_50_ value. The discrepancy between MPI5’s cellular inhibition potency and antiviral potency is likely due to two different cell lines used in two assays. Unlike 293T cells that are human cells, Vero E6 cells are African monkey cells. It is possible that MPI5 is more stable toward proteolytic digestion in Vero E6 cells than in 293T cells. Although the addition of CP-100356, a protyical ABC transporter inhibitor into growth media improved cellular potency of most MPI inhibitors but not to a dramatic level for most of them. Therefore, the main reason of low cellular and antiviral potency of MPI3 and other MPI inhibitors is not their active exportation from the inside of cells. Possible reasons that may contribute to low antiviral and cellular potency for these molecules include low cell permeability and proneness to both extracellular and intracellular proteolysis of these inhibitors. Although MPI8 is not the most potent MPI inhibitor according to its enzymatic inhibition potency, it has the best antiviral and cellular potency. The determined IC_50_ value using the new cellular assay for MPI8 is 31 nM that is less than a third of its enzymatic IC_50_ value. A likely reason is the possible accumulation of MPI8 in cells, which needs to be investigated. Other MPI inhibitors with high cellular potency are MPI5, 6, and 7. All display cellular M^Pro^ inhibition potency with an IC_50_ value below 1 μM. Among all 30 inhibitors we have tested, MPI5-8 show the most potency and warrant further investigation for possible use in COVID-19 treatment. As far as we know, MPI8 is the compound with the highest cellular M^Pro^ inhibition potency and the highest SARS-CoV-2 antiviral potency in Vero E6 cells. We recommend its urgent pharmacokinetic and pharmacodynamic characterization for COVID-19 clinical investigation.

GC376 is an investigational medication for treating feline infectious peritonitis, a lethal coronavirus disease in cats.^20^ Anivive Lifesciences Inc. is undergoing clinical investigation of repurposing GC376 for the treatment of COVID-19 patients. Although GC376 has high enzymatic M^Pro^ inhibition potency with an IC_50_ value of 30 nM, it shows relatively weak cellular M^Pro^ inhibition potency (IC_50_ > 2 μM). The cellular M^Pro^ inhibition potency of GC376 correlates with its antiviral potency that was determined with an EC_50_ value of 3.37 and 0.7 μM from two different studies.^17, 21^ In comparison to MPI8, GC376 is almost two orders of magnitude less potent in cellular M^Pro^ inhibition and in SARS-CoV-2 inhibition in Vero E6 cells. Low cellular permeability and stability likely contribute to this low cellular and antiviral potency. 11a is an M^Pro^ inhibitor that showed high antiviral potency with an EC_50_ value as 0.53 μM.^9^ However, its cellular M^Pro^ inhibition potency is much weaker in comparison to MPI5-8. Its estimated cellular IC_50_ value is higher than 2 μM. The discrepancy between cellular M^Pro^ inhibition potency and antiviral potency, although it is not dramatic, indicates that 11a may interfere with other critical process(es) in the SARS-CoV-2 life cycle to exert a potent antiviral effect, which needs to be explored.

Boceprevir and telaprevir are two medications approved for treating hepatitis C virus infection. Both have shown potency in inhibiting M^Pro^ enzymatically and boceprevir has also been characterized in an antiviral assay to show an EC_50_ value of 1.31 μM.^17^ However, both drugs display very weak potency in their cellular M^Pro^ inhibition tests. Since we detected very weak cellular inhibition of M^Pro^ for boceprevir at 10 μM, boceprevir must hit on other key step(s) in the SARS-CoV-2 pathogenesis and replication pathway to convene its high antiviral effect. Investigation in this possibility will likely lead to the discovery of novel target(s) for COVID-19 drug development. Other aldehyde and ketone-based inhibitors we have tested include calpeptin, MG-132, MG-115, calpain inhibitor II, and calpain inhibitor XII. Except calpain inhibitor XII that showed weak inhibition of M^Pro^ with an estimated IC_50_ value higher than 10 μM, all others exhibited close to undetectable cellular M^Pro^ inhibition up to 10 μM. Both calpain inhibitor II and XII have demonstrated antiviral potency toward SARS-CoV-2 with an EC_50_ value of 2.07 and 0.49 μM, respectively. Based on our cellular M^Pro^ inhibition analysis of two compounds, it is clear that their antiviral potency is not mainly from the inhibition of M^Pro^. Wang *et al*. have explored compounds with dual functions to inhibit both M^Pro^ and host calpains/cathepsins as antivirals for SARS-CoV-2.^32^ These compounds include calpain inhibitor II and XII. As such they likely inhibit host proteases to exert their potent antiviral effects. K777 weakly inhibited M^Pro^ in a kinetic assay but potent inhibited SARS-CoV-2 in an antiviral assay.^23^ It showed undetectable cellular M^Pro^ inhibition potency in our assay confirming that it must target other key process(es) in the SARS-CoV-2 life cycle.

Carmofur, tideglusib, ebselen, disulfiram, and PX-12 were discovered as M^Pro^ inhibitors from high-throughput screening. Although carmofur has an enzymatic IC_50_ value of 1.35 μM and generates a permanent covalent adduct with the M^Pro^ active site cysteine by forming a thiocarbamate, it showed undetectable cellular M^Pro^ inhibition potency up to 10 μM in our assay. This observation correlates well with its low antiviral potency.^24^ The high chemical reactivity of carmofur likely contributes to its low cellular and antiviral potency. Tideglusib, ebselen, disulfiram, and PX-12 are redox activity compounds that can form covalent adducts with the M^Pro^ active site cysteine. Except PX-12 that showed weak cellular potency at 10 μM, the other three drugs exhibited undetectable cellular M^Pro^ inhibition potency up to 10 μM. Among the four compounds, only ebselen has been examined in an antiviral assay.^7^ It has a determined EC_50_ value of 4.67 μM. Since ebselen showed undetectable cellular M^Pro^ inhibition up to 10 μM, its high antiviral potency must be from its interference with other key process(es) in the SARS-CoV-2 life cycle. The revelation of SARS-CoV-2 inhibition mechanism of ebselen will likely lead to the discovery of novel drug target(s) for COVID-19.

Bepridil is an antianginal drug with a demonstrated antiviral effect for SARS-CoV-2.^25^ It is an M^Pro^ inhibitor with an enzymatic IC_50_ value of 72 μM but a much lower antiviral EC_50_ value of 0.46 μM in ACE2^+^ A549 cells. Bepridil is known to inhibit other human viral pathogens as well.^33^ We detected close to undetectable cellular M^Pro^ inhibition potency for bepridil up to 10 μM. This correlates with its relatively high enzymatic IC_50_ value. Therefore, it is evident that bepridil must use a mechanism different from the inhibition of M^Pro^ in convening its high antiviral potency. This needs to be investigated. Chloroquine and hydroxychloroquine are two repurposed drugs for COVID-19 with demonstrated antiviral EC_50_ values as 5.47 and 0.72 μM, respectively.^34^ Although TMPRESS2 was shown as the target of chloroquine and hydroxychloroquine,^35^ a previous report showed that chloroquine and hydroxychloroquine potently inhibited M^Pro^ in an enzyme inhibition assay.^26^ We tested both drugs using the new cellular assay but revealed close to undetectable cellular M^Pro^ inhibition up to 10 μM for both drugs. We recharacterized enzymatic inhibition of M^Pro^ by both drugs. However, we were not able to detectable any M^Pro^ inhibition by hydroxychloroquine up to 16 μM and chloroquine exhibited weak inhibition of M^Pro^ at 16 μM. Based on our cellular data, enzymatic inhibition data, and data from a separate study,^36^ we are confident that both chloroquine and hydroxychloroquine don’t potently inhibit M^Pro^ inhibitors. Their antiviral activities are from different mechanism(s).

10-1, 10-2, and 10-3 are three diaryl esters in which 10-1 and 10-2 displayed high potency in inhibiting M^Pro^ enzymatically. All three compounds displayed significant cellular M^Pro^ inhibition potency at 10 μM but their potency is much lower than MPI5-8. Although 10-3 has much weaker enzymatic inhibition potency than 10-1 and 10-2, its cellular M^Pro^ inhibition potency is slightly higher than that from 10-1 and 10-2. A likely explanation is that 10-3 is more stable than 10-1 and 10-2 leading to a longer cellular time to convene its cellular M^Pro^ inhibition potency. Therefore, we recommend balancing cellular stability and enzymatic inhibition potency for future development of diaryl esters as M^Pro^ inhibitors to achieve optimal antiviral effects.

As a protyical ABC transporter inhibitor, CP-100356 can potentially improve intracellular accumulation of exogenous toxic molecules in cells. Providing CP-100356 improved cellular activity of all Opal inhibitors except MPI8. However, this improvement is limited. The highest improvement was observed for MPI7 that decreased the IC_50_ value from 0.19 μM to 0.075 μM. Since CP-100356 is not an approved medication, its use in combination with an M^Pro^ inhibitor for COVID-19 treatment will face significant hurdles in clearing out toxicity and other clinical concerns. Due to its nonsignificant improvement of cellular activity for an M^Pro^ inhibitor, we caution against its use. MPI8 showed similar cellular potency in the presence and absence of CP-100356, suggesting MPI8’s high propensity to accumulate inside cells that explains our observation that the determined cellular M^Pro^ inhibition IC_50_ value for MPI8 was threefold less than its determined enzymatic inhibition IC_50_ value. Data related to the use of CP-100356 supports that MPI8 is optimal for cellular M^Pro^ inhibition. As the compound with the highest cellular and antiviral potency among all literature and new compounds that we have tested in the current study, MPI8 is ready for further investigations in the treatment of COVID-19.

## CONCLUSION

We have developed a cellular assay for the determination of cellular potency of SARS-CoV-2 M^Pro^ inhibitors. Unlike an antiviral assay in which the interference of any key step in the SARS-CoV-2 life cycle may lead to a strong antiviral effect, this new cellular assay reveals only cellular M^Pro^ inhibition potency of a compound. It provides more precise information that reflects real M^Pro^ inhibition in cells than an antiviral assay. Using this assay, we characterized 30 M^Pro^ inhibitors. Our data indicated that 11a, boceprevir, ebselen, calpain inhibitor II, calpain inhibitor XII, K777, and bepridil likely interfere with key processes other than the M^Pro^ catalysis in the SARS-CoV-2 pathogenesis and replication pathways to convene their strong antiviral effects. Our results also revealed that MPI8 has the highest cellular potency among all compounds that were tested. It has a cellular M^Pro^ inhibition IC_50_ value of 31 nM. As the compound with the highest antiviral potency with an EC_50_ value of 30 nM, we cautiously believe and recommend that MPI8 is ready for preclinical and clinical investigations for COVID-19 treatment.

## MATERIALS AND METHODS

### Chemicals, reagents, and cell lines from commercial providers

We purchased HEK293T/17 cells from ATCC, DMEM with high glucose with GlutaMAX™ Supplement, fetal bovine serum, 0.25% Trypsin-EDTA, phenol red, puromycin, lipofectamine 3000, and dimethyl Sulfoxide from Thermo Fisher Scientific, linear polyethylenimine MW 25000 from Polysciences, RealTime-Glo™ annexin V apoptosis and necrosis assay kit from Promega, EndoFree plasmid DNA midi kit from Omega Bio-tek, antimycin a from Sigma Aldrich, GC376 from Selleck Chem, boceprevir, calpeptin, MG-132, telaprevir, and carmofur from MedChemExpress, ebselen from TCI, calpain inhibitors II and XII from Santa Cruz Biotechnology, MG-115 From Abcam, tideglusib, disulfiram and PX-12 from Cayman Chemical, chloroquine diphosphate from Alfa Aesar, hydroxychloroquine sulfate from Acros Organics, a fluorogenic M^Pro^ substrate DABCYL-Lys-Thr-Ser-Ala-Val-Leu-Gln-Ser-Gly-Phe-Arg-Lys-Met-Glu-EDANS from Bachem, and K777 as a gift from Prof. Thomas Meek at Texas A&M University. The synthesis of MPI1-9 and 11a were shown in a previous publication.^16^

### Plasmid construction

We amplified M^Pro^ with an *N*-terminal KTSAVLQ sequence using two primers FRET-M^pro^-for and FRET-M^pro^-rev primers (Table S1) and cloned it into the pECFP-18aa-EYFP plasmid (Addgene, #109330) between XhoI and HindIII restriction sites to afford pECFP-M^Pro^-EYFP. To construct pLVX-M^Pro^-eGFP-1, we amplified M^Pro^ with an *N*-terminal methionine using primers XbaI-Mpro-f and Mpro-HindIII-r (Table S1) and eGFP using primers HindIII-eGFP-f and eGFP-NotI-r. We digested the M^Pro^ fragment using XbaI and HindIII-HF restriction enzymes and the eGFP fragment using HindIII-HF and NotI restriction enzymes. We ligated the two digested fragments together with the pLVX-EF1α-IRES-Puro vector (Takara Bio 631988) that was digested at XbaI and NotI restriction sites. To facilitate the ligation of three fragments, we used a ratio of M^Pro^, eGFP and pLVX-EF1α-IRES-Puro digested products as 3:3:1. We constructed pLVX-M^Pro^-eGFP-2 in the same way as pLVX-M^Pro^-eGFP-1 except that we amplied the M^Pro^ fragment using primers XbaI-Cut-Mpro-f and Mpro-HindIII-r (Table S1). XbaI-Cut-Mpro-f encodes an MKTSAVLQ sequence for its integration to the M^Pro^ *N*-terminus.

### Transfection and MPI8 inhibition tests using pECFP-M^Pro^-EYFP

We grew 293T cells to 60% confluency and then transfected them with pECFP-M^Pro^-EYFP using Lipofectamine 3000. We added 10 μM MPI8 at the same time of transfection. After 72 h incubation, cells were collected and analyzed by flow cytometer as well as fluorescence microscopy. In order to obtain high-definition image, glass bottom plates were used for microimaging.

### Transfection and inhibition tests using pLVX-M^Pro^-eGFP-1 and pLVX-M^Pro^-eGFP-2

We grew 293T cells to 60% confluency and transfected them with pLVX-MPro-eGFP-1 or pLVX-using Lipofectamine 3000. We added Different concentration of MPI8 from nM to μM level at the same time of transfection. After 72 h incubation, we analyzed the transfected 293T cells using flow cytometry to determine fluorescent cell numbers and the eGFP fluorescent intensity.

### The establishment of 293T cells stably expressing M^Pro^-eGFP

To establish a 293T cell line that stably expresses M^Pro^-eGFP, we packaged lentivirus particles using the pLVX-M^Pro^-eGFP-2 plasmid. Briefly, we transfected 293T cells at 90% confluency with three plasmids including pLVX-M^Pro^-eGFP-2, pMD2.G and psPAX2 using 30 μg/mL polyethyleneimine. We collected supernatants at 48 h and 72 h after transfection separately. We concentrated and collected lentiviral particles from collected supernatant using Ultracentrifugation. We then transduced fresh 293T cells using the collected lentivirus particles. 48 h of transduction, we added puromycin the culture media to a final concentration of 2 μg/mL. We gradually raised the puromycin concentration 10 μg/mL in two weeks. The final stable cells were maintained in media containing 10 μg/mL puromycin.

### Apoptosis analysis

We performed the apoptosis analysis of the M^Pro^ stable cells and cells transiently transfected with the pLVX-M^Pro^-eGFP-2 plasmid using the RealTime-Glo™ Annexin V Apoptosis and Necrosis Assay kit from Promega. The cells were maintained in high glucose DMEM medium supplemented with 10% FBS, plated with a cell density of 5×10^5^ cells/ml. We set up five groups of experiments including 1) HEK 293T/17, 2) HEK 293T/17 + MPI8 (1 μM), 3) HEK 293T/17 cells stably expressing MPro-eGFP, **4)** HEK 293T/17 cells stably expressing MPro-eGFP + MPI8 (1 μM), **and 5)** HEK 293T/17 or HEK 293T/17 cells stably expressing MPro-eGFP + antimycin A (1 μM). Each experiment was repeated for 5 times. The assay was performed according to the instructor’s protocol. Chemiluminescence was recorded at 12, 24, 36, 48, 60, and 72 h after plating the cells. The luminescence readings were normalized using HEK 293T/17 as a negative control.

### Cellular M^Pro^ inhibition analysis for 29 selected compounds

We grew HEK 293T/17 cells in high-glucose DMEM with GlutaMAX Supplement and 10% fetal bovine serum in 10 cm culture plates under 37 □ and 5% CO_2_ to 80%~90% and then transfected cells with the pLVX-M^Pro^-eGFP-2 plasmid. For each transfection, we used 30 μg/mL polyethyleneimine and the total of 8 μg of the plasmid in 500 μL of the opti-MEM medium. We incubated cells with transfecting reagents for overnight. On the second day, we removed the medium, washed cells with a PBS buffer, digested them with 0.05% trypsin-EDTA, resuspended the cells in the original growth media, adjusted the cell density to 5 × 10^5^ cells/mL, provided 500 μL of suspended cells in the growth media to each well of a 48-well plate, and then added 100 μL of a drug solution in the growth media. These cells were then incubated under 37 □ and 5% CO_2_ for 72h before their flow cytometry analysis.

### Data collection, processing, and analysis

The cell was incubated with various concentrations of drugs in 37 °C for 3 days. After 3 days of incubation, we removed the media and then washed cells with 500 μL of PBS to remove dead cells. Cells were then trypsinized and spun down at 800 rpm for 5 min. We removed the supernatant and suspended the cell pellets in 200 μL of PBS. The fluorescence of each cell sample was collected by Cytoflex Beckman Flow Cytometer based on the size scatters (SSC-A and SSC-H) and forward scatter (FSC-A). We gated cells based on SSC-A and FSC-A then with SSC-A and SSC-H. The eGFP fluorescence was excited by blue laser (488 nm) and cells were collected at FITC-A (525 nm). After collecting the data, we analyzed and transferred data to csv files containing information of each cell sample. We then analyzed these files using a self-written MATLAB program for massive data processing. We sorted the FITC-A column from smallest to largest. A 10^6^ cutoff was set to separate the column to two groups, larger as positive and smaller as negative. We integrated the positive group and divided the total integrated fluorescent intensity by the total positive cell counts as Flu. Int. shown in all the graphs. The standard deviation of positive fluorescence was also calculated. It was then plotted and fitted non-linearly with an agonist curve (three parameters) against drug concentrations in the program Prism 9 (from GraphPAD for IC_50_ determination.

### Kinetic recharacterization of chloroquine and hydroxychloroquine

We prepared 10 mM stock solutions of hydroxychloroquinine (HCQ) and chloroquinine (CQ) in a PBS buffer and carried out IC_50_ assays for both HCQ and CQ by measuring activities of 50 nM M^pro^ against a concentration range of 0 to 16 μM HCQ and CQ. Serial dilutions of HCQ and CQ were carried out in the assay buffer by keeping the PBS concentration same. First, 100 nM M^pro^ in assay buffer (10 mM phosphate, 10 mM NaCl, 0.5 mM EDTA, pH 7.6) were treated with two times the working concentration of HCQ and CQ at 37 °C for 30 minutes. Then, 20 μM of the fluorogenic M^Pro^ substrate (prepared from 1 mM stock solution of the dye in DMSO) in the assay buffer was added to the reaction mixture to a final concentration of 10 μM. Immediately after the addition of the substrate, we started to monitor the reaction in a BioTek Neo2 plate reader with an excitation wavelength at 336 nM and emission detection at 490 nM. Initial product formation slopes at the first 5 minutes were calculated by simple linear regression and data were plotted in GraphPad Prism 9.

### The synthesis of 5-chloropyridin-3-yl 1H-indole-7-carboxylate (10-1)

To a solution of 5-chloropyridin-3-ol (1 mmol, 130 mg) and 1H-indole-7-carboxylic acid in anhydrous dichloromethane (DCM), we added DMAP (0.1 mmol, 12 mg) and EDC (1.2 mmol, 230 mg). The resulting solution was stirred at room temperature overnight. Then the reaction mixture was evaporated *in vacuo* and the residue was purified with flash chromatography to afford **10-1** as white solid (210 mg, 77%).

^1^H NMR (400 MHz, DMSO-*d*_6_) δ 11.34 (s, 1H), 8.65 (dd, *J* = 10.8, 2.2 Hz, 2H), 8.19 (t, *J* = 2.2 Hz, 1H), 8.03 – 7.92 (m, 2H), 7.47 (t, *J* = 2.9 Hz, 1H), 7.23 (t, *J* = 7.7 Hz, 1H), 6.65 (dd, *J* = 3.1, 1.9 Hz, 1H). ^13^C NMR (101 MHz, DMSO) δ 164.7, 147.9, 146.1, 143.0, 134.9, 131.2, 131.0, 130.2, 128.0, 127.9, 125.3, 119.2, 111.3, 102.5. ESI-HRMS: calculated for C_14_H_10_ClN_2_O_2_^+^: 273.0425; found: 273.0420.

### The synthesis of 5-chloropyridin-3-yl 1H-indole-4-carboxylate (10-2)

To a solution of 5-chloropyridin-3-ol (1 mmol, 130 mg) and 1H-indole-4-carboxylic acid in anhydrous DCM, we added DMAP (0.1 mmol, 12 mg) and EDC (1.2 mmol, 230 mg). The resulting solution was stirred at room temperature overnight. Then the reaction mixture was evaporated *in vacuo* and the residue was purified with flash chromatography to afford **10-2** as white solid (220 mg, 80%). ^1^H NMR (400 MHz, Chloroform-*d*) δ 8.72 (s, 1H), 8.53 (dd, *J* = 7.2, 2.2 Hz, 2H), 8.10 (dd, *J* = 7.5, 0.9 Hz, 1H), 7.75 (t, *J* = 2.2 Hz, 1H), 7.71 (dt, *J* = 8.1, 1.0 Hz, 1H), 7.43 (t, *J* = 2.9 Hz, 1H), 7.32 (t, *J* = 7.8 Hz, 1H), 7.23 (ddd, *J* = 3.2, 2.1, 0.9 Hz, 1H). ^13^C NMR (101 MHz, CDCl_3_) δ 165.0, 147.8, 145.7, 141.7, 136.7, 131.9, 130.0, 128.0, 127.2, 124.6, 121.3, 119.3, 117.4, 103.8. ESI-HRMS: calculated for C_14_H_10_ClN_2_O_2_^+^: 273.0425; found: 273.0420.

### The synthesis of 5-chloropyridin-3-yl 1H-indole-3-carboxylate (10-3)

To a solution of 5-chloropyridin-3-ol (1 mmol, 130 mg) and 1H-indole-3-carboxylic acid in anhydrous DCM, we added DMAP (0.1 mmol, 12 mg) and EDC (1.2 mmol, 230 mg). The resulting solution was stirred at room temperature overnight. Then the reaction mixture was evaporated *in vacuo* and the residue was purified with flash chromatography to afford **10-3** as white solid (190 mg, 69%). ^1^H NMR (400 MHz, DMSO-*d*_6_) δ 12.27 (s, 1H), 8.58 (dd, *J* = 2.3, 1.0 Hz, 2H), 8.40 (s, 1H), 8.08 (t, *J* = 2.2 Hz, 1H), 8.06 – 8.00 (m, 1H), 7.60 – 7.51 (m, 1H), 7.31 – 7.22 (m, 2H). ^13^C NMR (101 MHz, DMSO) δ 162.3, 148.0, 145.6, 142.8, 137.0, 135.1, 131.2, 130.8, 126.2, 123.4, 122.4, 120.8, 113.2, 104.8. ESI-HRMS: calculated for C_14_H_10_ClN_2_O_2_^+^: 273.0425; found: 273.0420.

### Kinetic characterization of 10-1, 10-2 and 10-3 in inhibiting M^Pro^

We performed M^Pro^ inhibition assays of these compounds using with the following assay buffer: 10 mM sodium phosphate, 10 mM NaCl and 0.5 mM EDTA in pH 7.6. We diluted a stock solution of the enzyme to 200 nM with the assay buffer. Stock solutions of inhibitors were prepared in DMSO. The fluorogenic M^Pro^ substrate was diluted to 20 μM in the assay buffer. The final concentrations in the enzymatic assay were 1.25 % DMSO, 2 μM DTT, 10 μM substrate and 20 nM M^Pro^. To perform the assays, we mixed 39 μL of the assay buffer, 1 μL inhibitor solution (or DMSO) and 10 μL of 200 nM M^Pro^ thoroughly and then incubated the solution at 37 °C for 30 min. The reaction was initiated by adding 50 μL of 20 μM substrate and the fluorescence intensity at 455 nm under 336 nm excitation was measured. We performed all experiments at ten different concentrations of three inhibitors in triplicate with both positive and negative controls. The initial rate was calculated according to the fluorescent intensity in the first five minutes by linear regression, which was then normalized according to the initial rate of positive and negative controls. IC_50_ curve was determined by Prism 9 from GraphPad.

### Characterization of cellular potency of MPI1-9, GC376, 11a, 10-1, 10-2, and 10-3 in the presence of CP-100356

All cellular M^Pro^ inhibition assays for these fourteen compounds were repeated with the addition of CP-100356 in DMSO to a final concentration of 0.5μM. The overall assay process and analysis were identical to the assays without CP-100356.

### Plaque reduction neutralization tests of SARS-CoV-2 by MPI5-8

We seeded 18 × 10^3^ Vero cells per well in flat bottom 96 well plates in a total volume of 200 uL of a culturing medium (DMEM + 10% FBS + glutamine) and incubated cells overnight at 37 °C and under 5 % CO_2_. Next day, we titrated compounds in separate round bottom 96 well plates using the culturing medium. We then discarded the original medium used for cell culturing and replaced it with 50 ul of compound-containing media from round bottom plates. We incubated cells for 2 h at 36 °C and under 5 % CO_2_. After incubation, we added 1000 PFU/50 uL of SARS-COV-2 (USA-WA1/2020) to each well and incubated at 36 °C and under 5% CO_2_ for 1 h. After incubation, we added 100ul of overlay (1:1 of 2% methylcellulose and the culture medium) was added to each well. We incubated plates for 3 days at 36 °C and under 5% CO_2_. Staining was performed by discarding the supernatant, fixing the plates with 4% paraformaldehyde in the PBS buffer for 30 minutes and staining with crystal violet. Plaques were then counted.

## ACKNOWLEDGEMENT

This work was supported by Welch Foundation (Grant A-1715 to W.R.Liu) and the Texas A&M University President’s Excellence Fund. We thank Prof. Thomas Meek for providing us the compound K777.

## AUTHOR CONTRIBUTIONS

W.R.L. conceived the project. W.C., C.-C.D.C., Z.Z.G., X.R.M., E.C.V., Y.R.A., Y.M., Y.Q., S.X., and W.R.L. designed and performed experiments. W.C, C.-C.D.C., Z.Z.G., S.X., and W.R.L. wrote the manuscript. All authors approved the final manuscript before submission.

