## supplementary information for "Cellular Activities of SARS-CoV-2 Main Protease Inhibitors Reveal Their Unique Characteristics"

**Supplementary Figures and Legends**

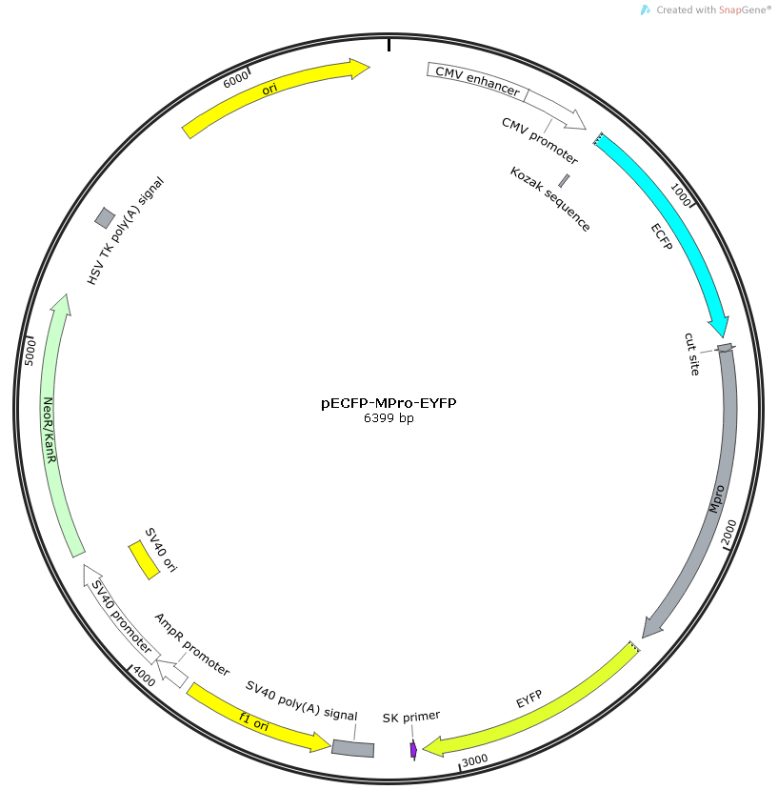

**Figure S1:** The plasmid map of pECFP-M^Pro^-EYFP

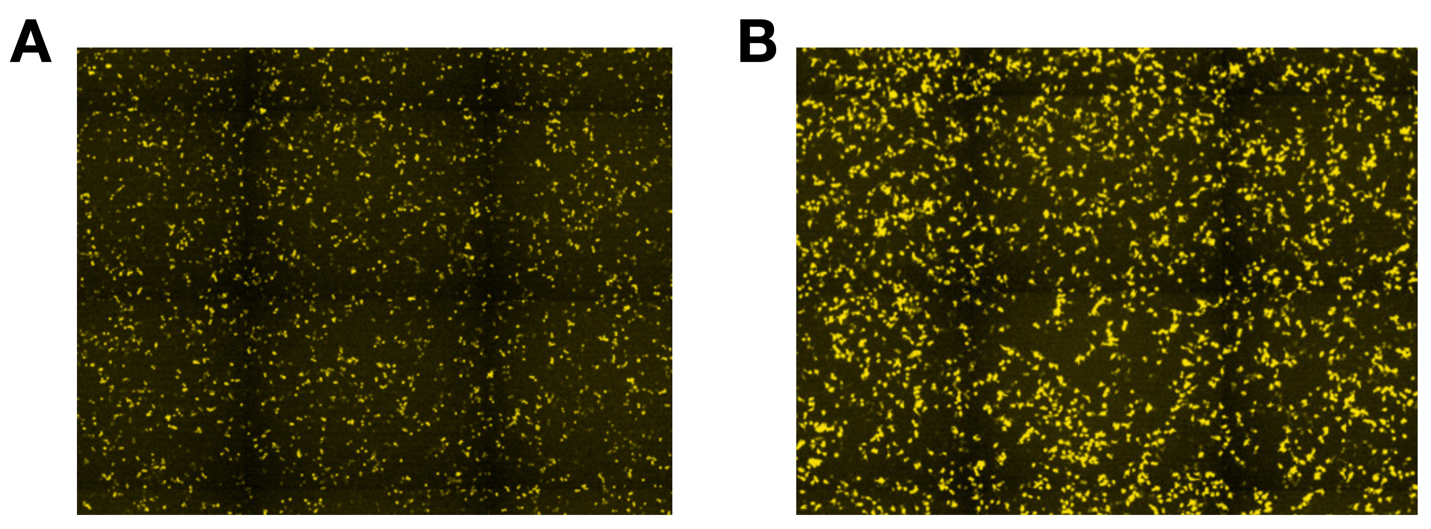

**Figure S2:** Yellow fluorescence from expressed CFP-M^Pro^-YFP in 293T cells transfected with pECFP-M^Pro^-EYFP and grown in the absence (**A**) or presence (**B**) of 10 μM MPI8.

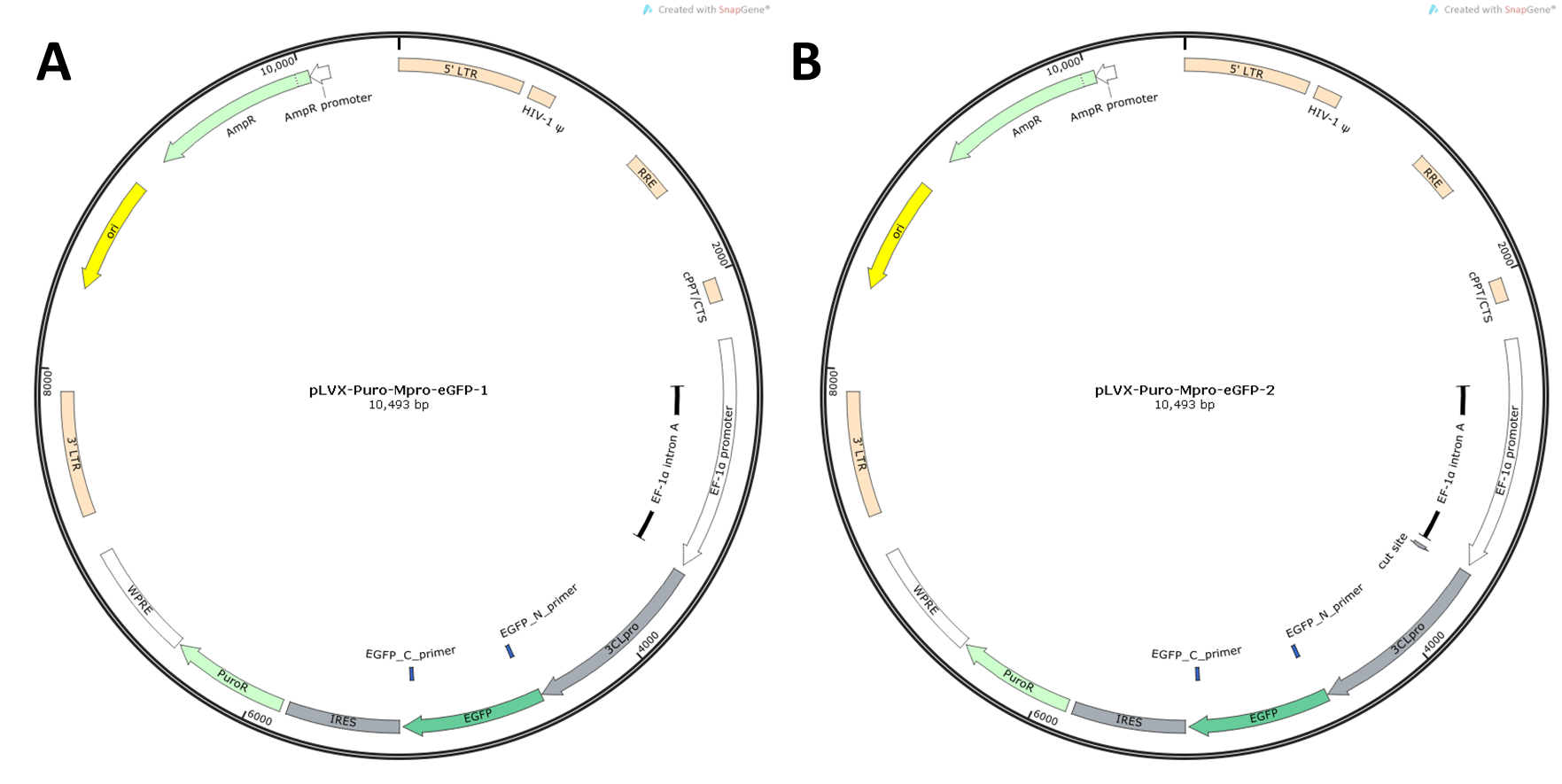

**Figure S3:** Plasmid maps of pLVX-Puro-M^Pro^-eGFP-1 (**A**) and pLVX-Puro-M^Pro^-eGFP-2 (**B**)

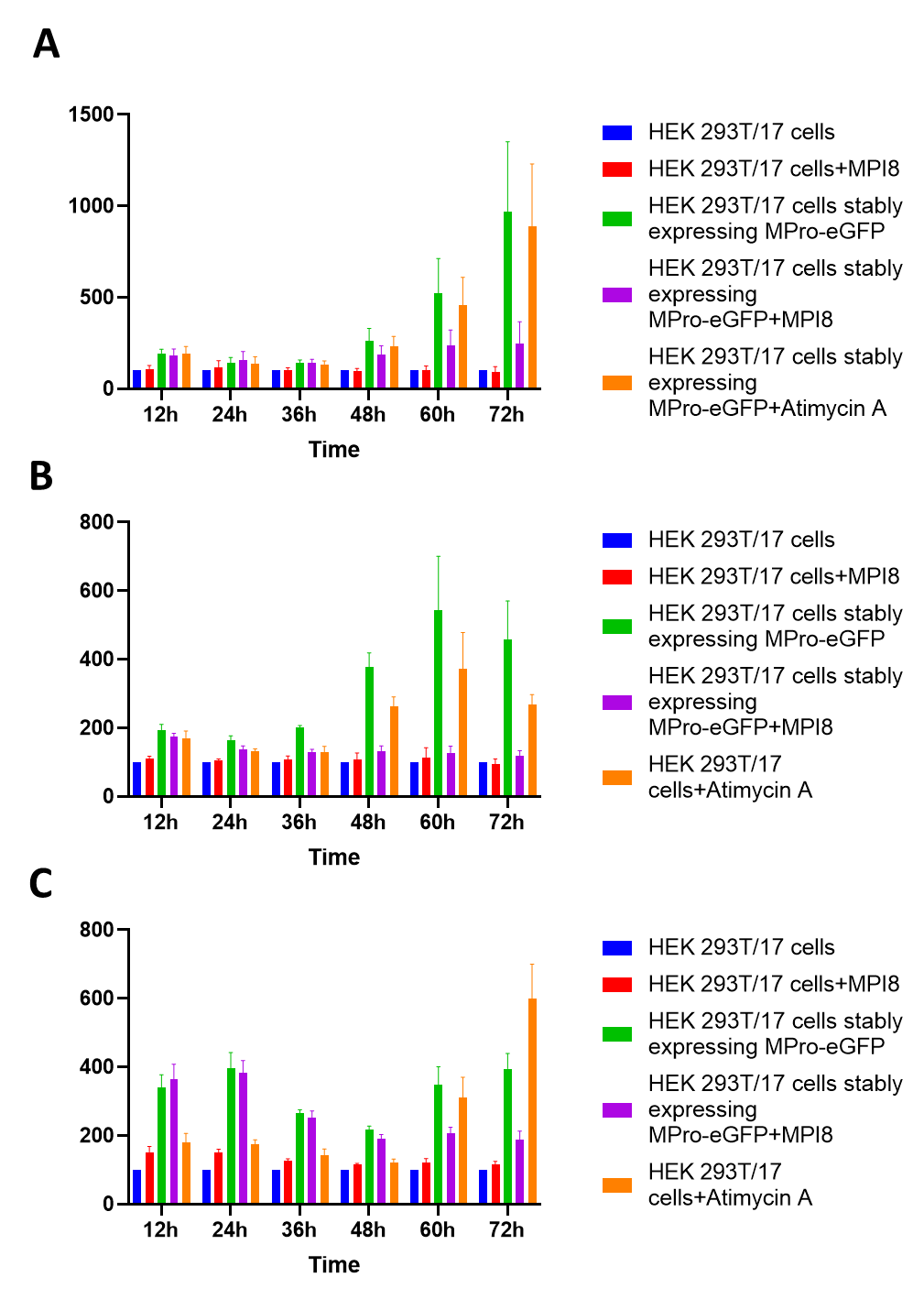

**Figure S4**: 293T/17 cells that were established in the presence of MPI8 exhibited strong apoptosis when MPI8 was withdrawn from the growth media. The cell assay was performed with RealTime-Glo™ Annexin V Apoptosis and Necrosis Assay kit from Promega. HEK 293T/17 and constructed HEK 293T/17 cells stably expressing M^Pro^-eGFP were used for this cell assay. The cells were maintained in high glucose DMEM medium supplemented with 10% FBS, plated with a cell density of 5×10^5^ cells/mL. Five groups of experiments were set:

HEK 293T/17;

HEK 293T/17 + MPI8 (1 μM)

HEK 293T/17 cells stably expressing M^Pro^-eGFP

HEK 293T/17 cells stably expressing M^Pro^-eGFP + MPI8 (1 μM)

HEK 293T/17(b&c) or HEK 293T/17 cells stably expressing M^Pro^-eGFP(a) + Antimycin A (1 μM)

Each experiment has 5 repeats.

The cell assay was performed as instructed by the protocol, luminescence was recorded at 12h, 24h, 36h, 48h, 60h, 72h after plating the cells. The luminescence readings were normalized using HEK 293T/17 as a negative control, which was set to a unit of 100.

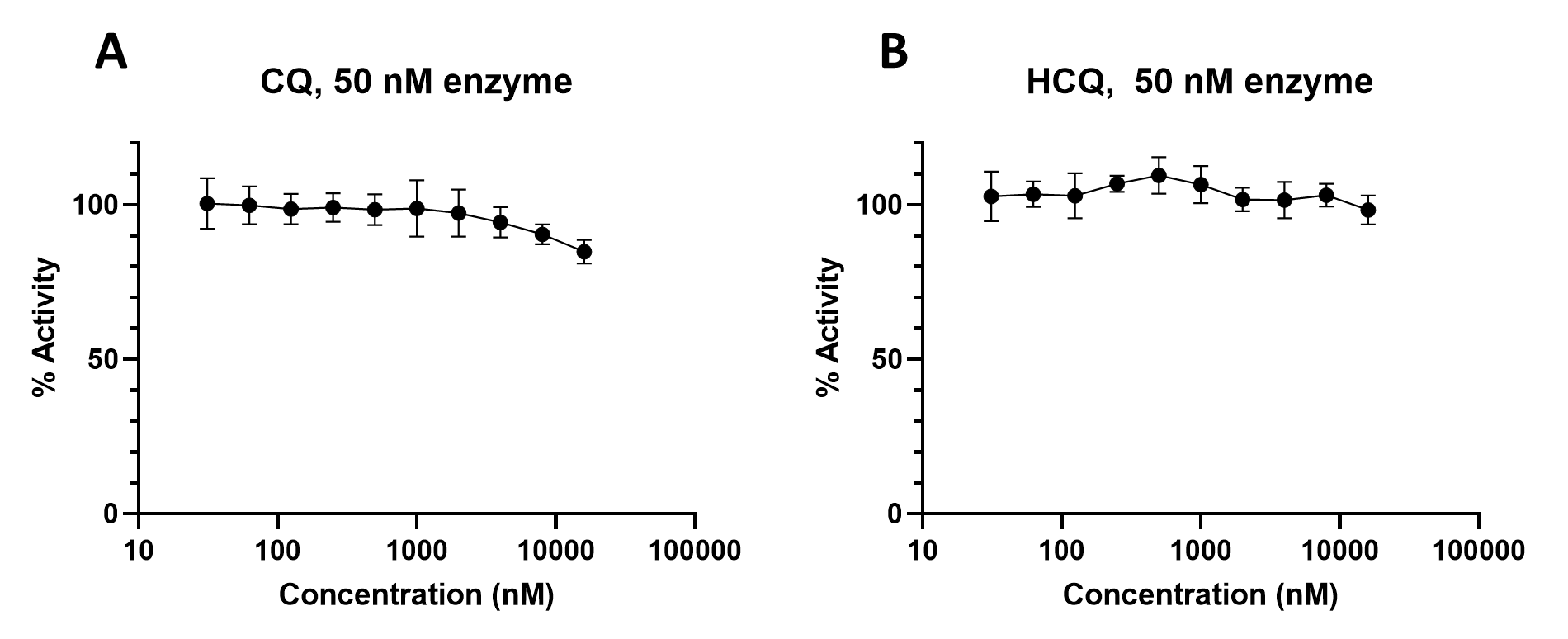

**Figure S5**: The recharacterization of M^Pro^ inhibition by (A) chloroquine and (B) hydroxychloroquine.

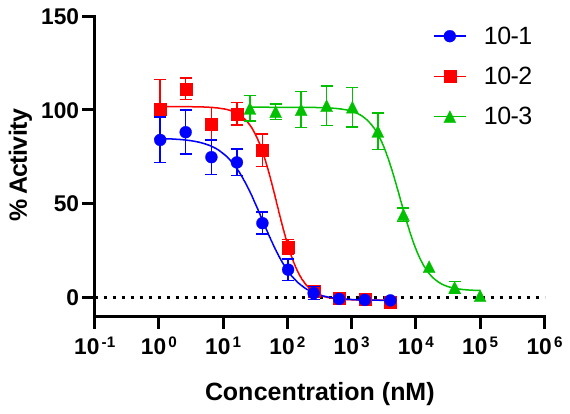

**Figure S6:** The kinetic characterization of 10-1, 10-2, and 10-3 in their inhibition of M^Pro^.

**Table S1:** The primers and their sequences used in the construction of plasmids.

| **Primer** | **Sequence** |
| --- | --- |
| FRET-Mpro-for | AGATCTCGAGTCAAAACAAGCGCGGTGC |
| FRET-Mpro-rev | TTCGAAGCTTGCTGAAAAGTTACGCCGGAAC |
| XbaI-Mpro-f | TAGTTCTAGAATGTCAGGGTTTCGCAAG |
| Mpro-HindIII-r | CCATAAGCTTGCCAAAAGTTACGCCGGAACAC |
| HinIII-eGFP-f | TGGCAAGCTTATGGTGAGCAAGGGC |
| eGFP-NotI-r | ATCCGCGGCCGCTTACTTGTACAGCTCGTCCATG |
| XbaI-Cut-Mpro-f | TAGTTCTAGAATGAAAACAAGCGCGGTGCTCCAGTCAGGGTTTCGCAAGATG |
